# Comparative study of GenePOC GBS LB Assay and GeneXpert GBS LB Assay for the detection of Group B *Streptococcus* in prenatal screening samples

**DOI:** 10.1101/553487

**Authors:** Tsokyi Choera, Brittney Jung-Hynes, Derrick J. Chen

## Abstract

Group B Streptococcal (GBS) infections in the United States are a leading cause of meningitis and sepsis in newborns. The CDC, therefore recommends GBS screening for all pregnant women at 35–37 weeks of gestation and administration of intrapartum prophylaxis (in those that tested positive) as an effective means of controlling disease transmission. Several FDA approved molecular diagnostic tests are available for rapid and accurate detection of GBS in antepartum women. In this study, we report a clinical comparison of the Xpert GBS LB assay and a novel FDA-cleared test, GenePOC GBS LB assay. A total of 250 vaginal-rectal swabs from women undergoing prenatal screening were submitted to the University of Wisconsin’s clinical microbiology laboratory for GBS testing. We found 96.8% of samples were concordant between the two tests, while 3.2% were discordant with higher sensitivity observed for the GenePOC GBS LB assay, and higher specificity for the GeneXpert GBS LB assay.

## Introduction

*Streptococcus agalactiae,* often referred to as Group B *Streptococcus* (GBS), is a gram-positive bacterium found in the rectum and vagina of approximately 25% of pregnant women (1). While GBS is an asymptomatic colonizer of most healthy adults, it can cause severe infections in neonates, including sepsis, pneumonia, and meningitis (2, 3). GBS early onset disease (EOD) are infections that occur in the first week of life and can be extremely dangerous to the newborn; EOD occurred in 3 per 1000 live births and was associated with a high mortality rate prior to the 1990s (4). Due to the high neonatal mortality rate caused by GBS infections, the Center for Disease Control (CDC) implemented a universal guideline in 1996 with recommended screening of all pregnant women at 35–37 weeks of gestation and administration of intrapartum prophylaxis (IAP) in pregnant women that tested positive (1, 5).

Despite these guidelines, infection with *S. agalactiae* remains a leading cause of morbidity in neonates born in the United States and therefore the implementation of more rapid and sensitive screening techniques for GBS detection may further reduce transmission of GBS infection intrapartum (1, 6, 7). The current gold standard for GBS detection is enrichment of the primary specimen, a vaginal-rectal swab, followed by subculture onto a blood agar plate with phenotypic characterization (1). This gold standard method has an extensive turnaround time and GBS colonization sensitivity is only 54-87% (8, 9). Furthermore, bacterial culture requires an experienced technician to further identify and test characteristics of GBS such agglutination and beta hemolysis (1).

Although culture remains the gold standard in GBS diagnostics, the 2010 CDC revision for GBS testing allows for nucleic acid amplification tests (NAAT) such as polymerase chain reaction (PCR) as an option for GBS testing (1). A PCR method based on amplification of the CAMP factor encoding gene (*cfb*), a fragment that is present in nearly all GBS strains, was developed by Ke et al. (3) in 2000. Since then, several rapid and sensitive DNA probes and NAAT assays for GBS have been developed and approved or cleared by the Food and Drug Administration (FDA) (3, 10, 11). In our study, we report a clinical comparison of two FDA cleared NAATs for the detection of GBS in antepartum women: Xpert^®^ GBS LB (Cepheid Inc., Sunnyvale, CA, USA) and a novel recently FDA-cleared test, GenePOC^™^ GBS LB (GenePOC inc, Québec City, Canada). Both systems perform an automated nucleic acid extraction, real-time PCR amplification, and detection of the target nucleic acid sequences after LIM broth enrichment (12, 13).

## Materials and Methods

### Study specimens

A total of 250 vaginal-rectal swabs from pregnant women undergoing routine GBS screening were tested in the present study. The specimens were submitted to the University of Wisconsin’s clinical microbiology laboratory between December 2017 to February 2018. All swab specimens were enriched in Lim broth for 18-24 hours. Post-enrichment, each sample was clinically tested using Xpert^®^ GBS LB assay, followed by testing with GenePOC^™^ GBS LB assay within 72 hours. All results were recorded and repeat testing on each of the instrument platforms and cultures were performed for any discordant specimens. The same positive (ATCC 12386, *Streptococcus agalactiae*) and negative (ATCC 9809, *Streptococcus gallolyticus*) external controls were used in both assays.

### Xpert^®^ GBS LB Assay

All enriched samples were first tested on the GeneXpert Dx system using the Xpert® GBS LB Assay according to the manufacturer’s instructions. Briefly, the LIM broth enriched samples were inverted 3 times and a sterile swab was immersed into the LIM broth enriched samples. This swab was immediately inserted into the sample chamber in the Xpert^®^ GBS LB cartridge and run on the GeneXpert Dx System.

### GenePOC™ GBS LB Assay

Following completion on the GeneXpert Dx system, the same LIM broth enriched specimens were tested on the GenePOC™ system according to the manufacturer’s instructions. Briefly, the LIM broth enriched samples were vortexed for 15 seconds and diluted into the diluent buffer. The diluted samples were vortexed for another 15 seconds and loaded into GenePOC™ GBS LB PIE and run on the revogene.

### Bacterial Culture Testing

Reference culture testing was performed on all discordant samples in accordance with published CDC guidelines. Briefly, the LIM broth enriched samples were sub-cultured onto blood agar plates and incubated for 18 to 24h at 37°C in 5% CO_2_. The blood agar plates without any colonies observed at 24h were re-incubated for an additional 24h. If after 48h no colonies formed, the culture was deemed negative. Any suspected colonies were identified via matrix-assisted laser desorption ionization time-of-flight mass spectrometry (MALDI-TOF MS). For MALDI-TOF MS-identified *Streptococcus agalactiae* isolates, 16s rRNA gene sequencing was conducted for confirmation.

### Time and cost

The hands-on setup time of each assay was timed and averaged over five individual runs. List price of each test and instrument were provided by vendors.

### Data analysis

Results from both GeneXpert^®^ and GenePOC^™^ systems were compared for accuracy. Concordant results between the assays were recorded, and no additional testing was performed. Discordant results between the initial tests were resolved in two ways: repeating the tests on both instrument platforms and culturing the enriched specimens to check for GBS presence. Results from these additional tests were compared against two definitions of true positivity: 3 of 4 molecular tests yielding a positive GBS result or GBS presence in culture. These definitions will be referred to as standard A and B, respectively.

## Results

From the 250 total samples tested, 242 samples were concordant (96.8%), and eight samples were discrepant (3.2%) between the two molecular assays. Among the eight discrepant results, seven specimens tested negative by GeneXpert^®^, but positive by GenePOC^™^ and one tested positive by GeneXpert^®^, but negative by GenePOC^™^ with initial testing. All discrepant results were repeated by both molecular assays and set up for culture. Upon repeat testing, one discrepant sample that initially tested positive by GeneXpert^®^, tested negative by both standard A and standard B definitions. Upon repeat of the seven discrepant samples that initially tested negative by GeneXpert^®^, three samples tested positive and three samples tested negative by standard A; and these six samples tested negative by standard B. Interestingly, one of the samples that had initially tested negative on the GeneXpert^®^ tested negative upon repeat on GeneXpert^®^ but tested positive on GenePOC^™^ (initial and repeat) and had growth in culture. The strain isolated from that specimen was confirmed to be *Streptococcus agalactiae* by sequencing the entire 16S rRNA gene and by MALDI-TOF MS analysis. The results are summarized in Table 1A. For all discrepant results, Ct value (listed in Table 1B) were obtained on repeat test from either the instrument (GeneXpert^®^) or by contacting the vendor (GenePOC^™^).

**Table 1A.** Comparison of the results of Xpert GBS LB Assay, GenePOC GBS LB Assay, and Culture

| No. of specimens | Test Results |  |  |  |  | Results based on true positive definitions |  |
| --- | --- | --- | --- | --- | --- | --- | --- |
|  | Xpert GBS LB assay | GenePOC GBS LB assay | Repeat Xpert | Repeat GenePOC | Culture | 3 of 4 Positive Molecular Tests | Culture Positive |
| 192 | Negative | Negative | N/A | N/A | N/A | N/A | N/A |
| 50 | Positive | Positive | N/A | N/A | N/A | N/A | N/A |
| 3 | Negative | Positive | Positive | Positive | Negative | Positive | Negative |
| 2 | Negative | Positive | Negative | Negative | Negative | Negative | Negative |
| 1 | Positive | Negative | Negative | Negative | Negative | Negative | Negative |
| 1 | Negative | Positive | Negative | Positive | Negative | Negative | Negative |
| 1 | Negative | Positive | Negative | Positive | Positive | Negative | Positive |

**Table 1B.** Results of Discrepant Specimens

| Discrepant Specimen | Xpert GBS LB assay | GenePOC GBS LB assay | Repeat Xpert | Repeat Xpert Ct value | Repeat GenePOC | Repeat GenePOC Ct value | Culture |
| --- | --- | --- | --- | --- | --- | --- | --- |
| 1 | Negative | Positive | Positive | 40.2 | Positive | 39 | Negative |
| 2 | Negative | Positive | Positive | 37.7 | Positive | 39 | Negative |
| 3 | Negative | Positive | Positive | 33.4 | Positive | 36 | Negative |
| 4 | Negative | Positive | Negative | 42.8 | Negative | 48 | Negative |
| 5 | Negative | Positive | Negative | ND* | Negative | 48 | Negative |
| 6 | Positive | Negative | Negative | ND* | Negative | ND* | Negative |
| 7 | Negative | Positive | Negative | 38.4 | Positive | 48 | Negative |
| 8 | Negative | Positive | Negative | ND* | Positive | 23 | Positive |

\*ND: Not Detected

Sensitivity and specificity of each of the assays were assessed according to the two standards of true positivity and the results are displayed in Table 2. When standard A was applied to define true positives for the detections of GBS, GenePOC^™^ demonstrated a sensitivity and specificity of 100% and 98%, respectively, while GeneXpert^®^ demonstrated a sensitivity and specificity of 94% and 99%, respectively. When standard B was applied, GenePOC^™^ demonstrated a sensitivity and specificity of 100% and 97%, respectively, while GeneXpert^®^ demonstrated a sensitivity and specificity of 98% and 100%, respectively.

**Table 2.** Sensitivity and Specificity

| Definition of Positivity | 3 of 4 Molecular Tests |  | Culture |  |
| --- | --- | --- | --- | --- |
| Assay | % sensitivity | % specificity | % sensitivity | % specificity |
| Xpert GBS LB | 94% | 99% | 98% | 100% |
| Gene POC GBS LB | 100% | 98% | 100% | 97% |

Time and cost of each assay were also assessed as displayed in Table 3. The GenePOC^™^ GBS LB test resulted in an average of 2.7 minutes to setup with a run time of 70 mins, while the Xpert GBS LB assay resulted in an average of 1.3 minutes to setup with a run time of 59 min per test. The list price of each test was $28 for GenePOC^™^ and $30 for GeneXpert^®^. The list price of the instrument holding an equal number of testing modules was $35,000 for GenePOC^™^ and $110,000 for GeneXpert^®^.

**Table 3.** Time and Material Cost/Test for GBS Assay

| Assay | Time (mins) | | Cost (\$)* | |
| --- | --- | --- | --- | --- |
|  | Setup Per Test | Run Time | Per Test* | Per Instrument* |
| Xpert GBS LB | 1.3 | 59 | 30 | 110,000 |
| Gene POC GBS LB | 2.7 | 70 | 28 | 35,000 |

\*Cost is based on the list price provided by the vendors, however each vendor works with individual labs to try to make these kits and instrument affordable within their budgets
\*\*The test price is listed as per test; however the tests are purchased as kits

## Discussion

The purpose of our study was to compare the newly FDA-cleared GenePOC^™^ GBS LB assay to the GeneXpert^®^ GBS LB assay. Whereas, the small consumable and equipment size of GenePOC^™^ are a significant advantage as it translates to minimum space requirements and better waste management, GeneXpert^®^ allows the flexibility of adding on more modules, has a more extensive test menu, and specimens are tested individually; the GenePOC^™^ instrument allows batched testing of 1 to 8 samples per run (12, 13). One shortcoming of the GenePOC^™^ instrument during our testing was that we were not notified of any error when loading the pie onto the instrument until run was complete; this caused error/invalid rates of 2.4% for GenePOC^™^, while GeneXpert^®^ had a 0% error/invalid rate. An additional challenge of GenePOC^™^ instrument during this study was that we were unable to readily view Ct values, which could have been useful for troubleshooting and discrepancy analysis; we were notified that this feature will be made available in the future. A major advantage of GenePOC^™^ was the higher sensitivity, although there were more false positives based on our two standards of true positivity. It is noteworthy to highlight the single discrepant sample that resulted in GBS-negative GeneXpert^®^ results compared to GBS-positive GenePOC^™^ and culture results (Table 1A). Both the GeneXpert^®^ and the GenePOC^™^ GBS assays’ primers and probes detect a target within or adjacent to the CAMP factor encoding gene (*cfb*) of *S. agalactiae* (4, 12, 13). This *cfb* gene is noted to be present in almost all group B streptococci, yet recent findings have described cases of *cfb*-negative isolates that were missed by the GeneXpert^®^ system (15, 16). It is likely that this isolate is either missing or has a mutation in the primer binding region for the primer pair of the GeneXpert^®^, and GenePOC^™^ GBS assay’s primers and probe amplify a *cfb* site that is different from that of GeneXpert^®^. Sequencing of the *cfb* gene and adjacent sites of this strain may provide further clarification on why this strain was missed by the Xpert assay. The GeneXpert^®^ false negative GBS strain raises an alarming concern and will need to be explored in future studies.

To summarize, we observed higher sensitivity for the GenePOC^™^ assay, and higher specificity for the GeneXpert^®^ assay when applying the two standards of true positivity. Both the setup and run times were slightly longer on the GenePOC^™^ assay compared to that of the GeneXpert^®^ assay. The extra dilution and vortexing step added additional time and material, such as variable volume pipettes to the setup of the GenePOC^™^ GBS LB assay, in comparison to the Xpert^®^ GBS LB assay. The list price per test was less for the GenePOC^™^ assay compared to that of GeneXpert^®^ assay. For the same number of testing modules, the GenePOC^™^ instrument costs less than the GeneXpert^®^ Dx System based on list prices. Overall, both assays perform well and agree in results, however there were a few discrepancies and specific differences listed in our study between the platforms that labs should consider when deciding between the two platforms and GBS assays.

## Acknowledgment

GenePOC provided the revogene instrument and GenePOC GBS LB reagents used in this study.

